# Drug screen reveals new potent host-targeted antivirals against Mpox virus

**DOI:** 10.1101/2025.05.02.651913

**Authors:** Arjit Vijey Jeyachandran, Anne K. Zaiss, Nikhil Chakravarty, Sneha Singh, Yennifer Delgado, Ramya Paravastu, Nivedha Satheeshkumar, Ephrem Gerald, Aakash Jeysankar, Joshua Thomas, Lilly Fuller, Noella Lee, Cameron Taylor, Shantanu Joshi, Mark Parcells, Samuel W. French, Abhijit Date, Mehdi Bouhaddou, Gustavo Garcia, Ashok Kumar, Robert Damoiseaux, Vaithilingaraja Arumugaswami

## Abstract

Mpox virus (MPXV), a re-emerging zoonotic threat, has caused outbreaks in non-endemic regions through respiratory, sexual, and close-contact transmission. The increased transmissibility of Clade IIb fueled the 2022 global outbreak, with 2024 Clade Ib spread in the Democratic Republic of Congo further escalating concern. Both outbreaks were declared public health emergencies by the WHO. Although tecovirimat (TPOXX) has been used off-label for Mpox, its limited effectiveness highlights the critical need for newer antivirals for MPXV. We conducted high-throughput antiviral drug screening using a host-directed kinase inhibitor library composed of 2,750 compounds against 2022 Clade IIb MPXV. Our primary screen identified 138 compounds preventing MPXV cytopathic effects, including multiple inhibitors of EGFR, PI3K-mTOR, and Ras/Raf, as well as apoptosis and autophagy regulators. Secondary and tertiary screenings yielded a shortlist of potent, nontoxic antiviral compounds that inhibited MPXV replication. Three selected compounds, IRAK4-IN-6, SM-7368, and KRAS inhibitor-10, reduced MPXV-induced cell death in primary human epidermal keratinocytes. IRAK4-IN-6 and SM-7368 were also found to modulate NF-κB and STING signaling. Furthermore, these compounds were found effective in reducing skin lesions and viral burden in a mouse model of MPXV skin infection. Together, our study reveals new classes of antiviral compounds against MPXV, offering promising candidates for future clinical development.

## INTRODUCTION

Mpox virus (MPXV), formerly known as Monkeypox virus, was the causative agent of a global outbreak and public health emergency of international concern in 2022, which resulted in over 100,000 infections and 250 deaths in 122 countries, including 115 countries with no previously reported cases of Mpox (monkeypox disease)^1,2^. MPXV is an enveloped, double-stranded DNA virus (genus *Orthopoxviridae*) typically transmitted zoonotically from primates and rodents to humans^3^. However, recent MPXV outbreaks have been distinguished by sustained human-to-human transmission^3^, highlighting a key shift in the epidemiology of Mpox. MPXV primarily spreads through respiratory droplets and direct contact with exposed skin and mucosal surfaces^4^. Following an initial incubation period of 5-21 days, Mpox presents with fever, severe headache, lymphadenopathy, and muscle aches, along with characteristic rashes and lesions on the face and extremities^5,6^, and ocular manifestations, including eyelid papules, conjunctivitis, cataracts, and corneal erosion^7^, lasting approximately two weeks, and, sometimes, death. Given its 96.3% genetic similarity to the smallpox virus, which caused an estimated 300–500 million deaths in the 20th century, the emergence of Mpox poses a significant global health concern^8,9^. The current standard of care for Mpox includes the use of prophylaxis with the JYNNEOS smallpox vaccine (attenuated varicella virus (VACV) strain)^8,10^, which has shown 75% protection against MPXV infection^11^. The primary approach for Mpox symptom management is supportive therapy, such as wound care, pain relief, and fever reduction^12^.

Though primarily endemic to West and Central Africa^13^, distinct clades of MPXV have become responsible for viral spread in different world regions. Clade Ia was first isolated in the Congo Basin and shows mostly zoonotic spread^13,14^. Mainly endemic to Central Africa, Clade Ia has shown significant similarity (99% nucleotide sequence similarity) to West African Clade II^13^. Both clades are associated with severe disease, resulting in mortality. Clade Ia has a higher level of mortality (10%) as compared to Clade II (1%)^15,16^. The 2003 Mpox outbreak in the United States caused by Clade IIa presented the first worldwide spread of this virus, though the spread was still primarily zoonotic^17^. The 2022 global outbreak caused by Clade IIb demonstrated much higher human-to-human transmissibility than Clade Ia or IIa^18^. The newly emergent Clade Ib was recently identified as a causative agent of ongoing outbreaks in the Democratic Republic of Congo (DRC) and once again resulted in Mpox spread being declared a public health emergency of international concern by the World Health Organization^19^. This outbreak has caused nearly 2,500 infections across bordering African nations as of January 2025^20^, a count that is likely higher due to limitations in diagnostic and reporting infrastructure. Travel-associated Clade Ib cases have also been reported in India, the United States, and Europe^2^. Reports of Clade Ib in the DRC have shown higher rates of sexual transmission compared to other strains, disproportionately affecting women and children^21,22^. What has set Clade Ib MPXV apart from other clades is the rapid evolution of this strain, as well as the significantly increased virulence and transmissibility^23,24^.

Unlike other DNA viruses, which typically replicate in the nucleus of infected cells, poxviruses replicate in the cytosol^25^. MPXV utilizes cellular machinery and signaling pathways to complete its viral life cycle^25^. Additionally, MPXV can evade innate immune activity triggered by the presence of cytosolic DNA via the cGMP-AMP synthase (cGAS)-Stimulator of Interferon Response cGAMP Interactor 1 (STING) pathway^26^. The cGAS-STING pathway utilizes various kinases and transcription factors to activate interferons (IFNs), including IFN-β, to mount an innate immune response against viral infection^27^. Poxviruses encode immunomodulatory proteins responsible for evading innate immunity through methods including virostealth, virotransduction, and viromimicry^6,28^.

There are no currently approved therapeutic agents specific to Mpox. Tecovirimat (TPOXX), an agent initially developed to inhibit the smallpox VP37 viral protein^29^, is currently used to treat Mpox lesions as an off-label medication^30^. By inhibiting VP37 protein synthesis, TPOXX blocks the maturation and budding of viral particles from the cells, thereby hampering virus release from the cell. However, tecovirimat resistance has been reported in severely immunocompromised Mpox patients with a history of tecovirimat use^31^. Additionally, the landmark STOMP clinical study of 7,100 Mpox patients reported in November 2024 revealed that tecovirimat did not reduce the time to resolution of Mpox-related lesions and presented no definitive benefit^30^. Given the urgent and unmet need for effective antiviral therapies for Mpox, particularly in light of tecovirimat’s limited clinical efficacy and emerging resistance, we pursued a novel strategy to identify host-directed compounds with antiviral activity against MPXV, evaluating their efficacy both *in vitro* with relevant epithelial cell systems and *in vivo* through the establishment of a dermal mouse model of Mpox.

## RESULTS

### High-Throughput Screening of Antivirals against MPXV

Viruses rely on the host cell machinery for their replication and have evolved methods to modulate and hijack host cell signaling pathways, circumvent cell cycle checkpoints, influence cell proliferation, and evade immune responses^26,32^. Given the potential for rapid deployment, we selected a well-curated kinase inhibitor library at UCLA’s Molecular Screening Shared Resource, consisting of 2,750 small molecules designed to target host kinases to evaluate their ability to inhibit the 2022 Clade IIb MPXV strain (Fig. 1A). Some of these compounds have been investigated in human clinical trials. This approach leverages the central role of kinases in regulating key cellular processes and signaling pathways, making them attractive targets for uncovering host factors critical to viral replication and identifying candidates for antiviral development.

**Figure 1.**
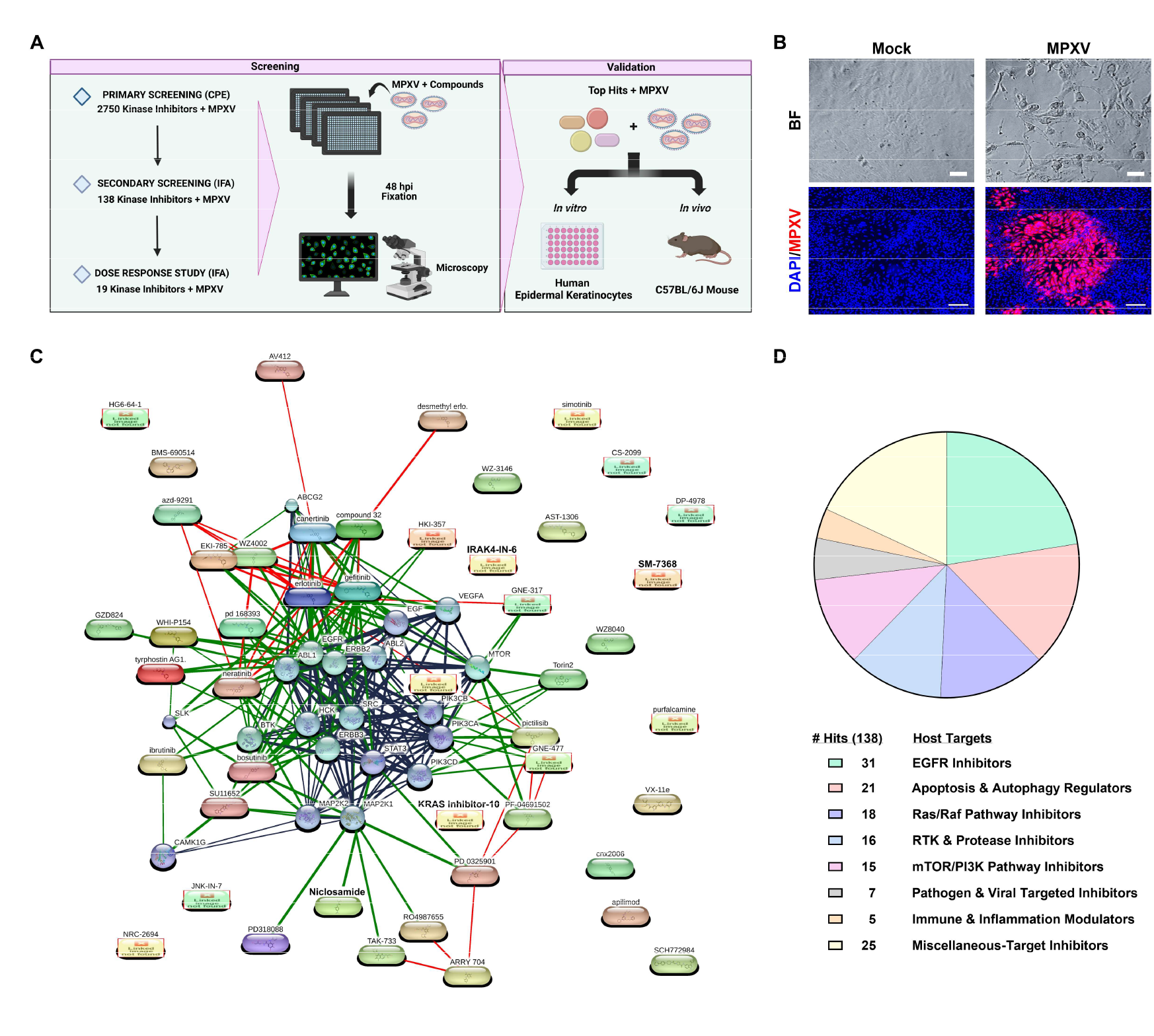
High-Throughput Primary Screening of Kinase Inhibitor Library Unveils Antiviral Compounds against MPXV. **(A)** Schematic representation of the high-throughput antiviral screening process. **(B)** Representative brightfield (BF) microscopy images of ARPE19 cells show the cytopathic effect induced by MPXV infection (MOI 0.1) at 72-hour post-infection as a part of (CPE)-based screening assay. Immunofluorescence (IF) images display MPXV A4L viral antigen (red) and DAPI-stained nuclei (blue), highlighting the extent of viral infection. The IF images were used for identifying anti-MPXV compounds as part of the high-throughput antiviral screening assay. Scale bar: 100 µm (BF), 200 µm (IF). **(C)** The connectivity map depicts host targets and their corresponding drug hits identified in the primary screen. The confidence-based network uses edge thickness to indicate the strength of the associations to represent protein-protein interaction (blue), chemical-protein interactions (green), and chemical-chemical interactions (red). Circular nodes represent proteins while oval nodes indicate compounds identified in the primary screen. **(D)** Pie chart depicting the distribution of host targets associated with drug molecules identified in the primary screen. The chart represents the allocation of 138 identified drug hits across key target pathways, as well as the number of hits corresponding to each pathway.

To establish this screening platform, we first evaluated the antiviral activity of tecovirimat, approved for smallpox. However, our preliminary data showed that tecovirimat did not exhibit strong antiviral activity against MPXV Clade IIb, as demonstrated by immunofluorescence analysis (Fig. S1). This data further supports the findings of the recent STOMP clinical trial, showing the reduced efficacy of tecovirimat in treating Mpox lesions in patients^33^ and emphasizing the need for newer antiviral strategies for MPXV.

In addition to typical skin rashes, MPXV causes moderate to severe ophthalmic manifestations^7^. Thus, we utilized the human ocular epithelial cell line ARPE-19 for the primary drug screen. Cells were seeded in 384-well plates and pretreated with each drug at a 1.6µM concentration for 16 hours. This lower dosage was selected to ensure that potent drug candidates would fall within the therapeutic range suitable for human use, as inhibiting kinases can be deleterious to cell health. The next day, the cells were infected with MPXV strain hMPXV/USA/MA001/2022 (Lineage B.1, Clade IIb) at MOI 0.1 and, at 72 hours post-infection (hpi), cytopathic effects (CPE), characterized by the fragmentation and rounding of cells, were evaluated using brightfield microscopy (Fig. 1B). This initial screen led to the identification of 138 compounds that inhibited MPXV-mediated CPE (Table S1). Chemical-protein interaction network mapping revealed that the identified hits clustered around key signaling pathways, including EGF, MAP2K, PI3K-mTOR, Ras/Raf, and NF-κB (Fig. 1C). Notably, 31 drug candidates were EGFR inhibitors (Fig. 1D). This was followed by 21 regulating apoptosis and autophagy, 18 inhibiting the Ras/Raf pathway, 16 inhibiting Receptor Tyrosine Kinase (RTK) and proteases, and 15 mTOR/PI3K pathway inhibitors. Additionally, 25 hits were broad-spectrum and multi-target inhibitors, highlighting their potential to modulate multiple pathways simultaneously (Fig. 1D). These findings suggest that MPXV-mediated cell injury processes, as well as viral replication, may be directed through these cellular pathways, uncovering potential targets for therapeutic interventions.

Next, all 138 compounds were subjected to a secondary screen at three concentrations with ten-fold dilutions (1.66 µM, 0.16 µM, and 0.016 µM) in ARPE-19 cells. Like the primary screen, cells were pretreated with each compound and, 16 hours later, were infected with MPXV at MOI 0.1 and immunostained with an MPXV-A4L antibody at 48hpi. Automated confocal microscopy imaging identified 19 compounds that reduced the number of MPXV-infected cells, indicating ~80% inhibition of viral replication compared to the vehicle-treated control at 1.66µM (Fig. 2A, 2B). Among the 19 compounds, seven (GZD824, GSK2126458, Pelitinib, SM-7368, INK-128, Mefatinib, and PI3K/mTOR inhibitor-11) showed potent antiviral activity even at lower concentrations (0.16 µM and 0.016 µM) (Fig. S1). The 19 kinase inhibitors that actively inhibited MPXV replication were clustered into the following classes: seven mTOR pathway regulators, four RTK and Non-Receptor Tyrosine Kinase (NRTK) inhibitors, three immune and inflammation modulators, two EGFR inhibitors, two Ras/Raf pathway inhibitors, and one helminthic DNA-targeting drug (Fig. 2C).

**Figure 2.**
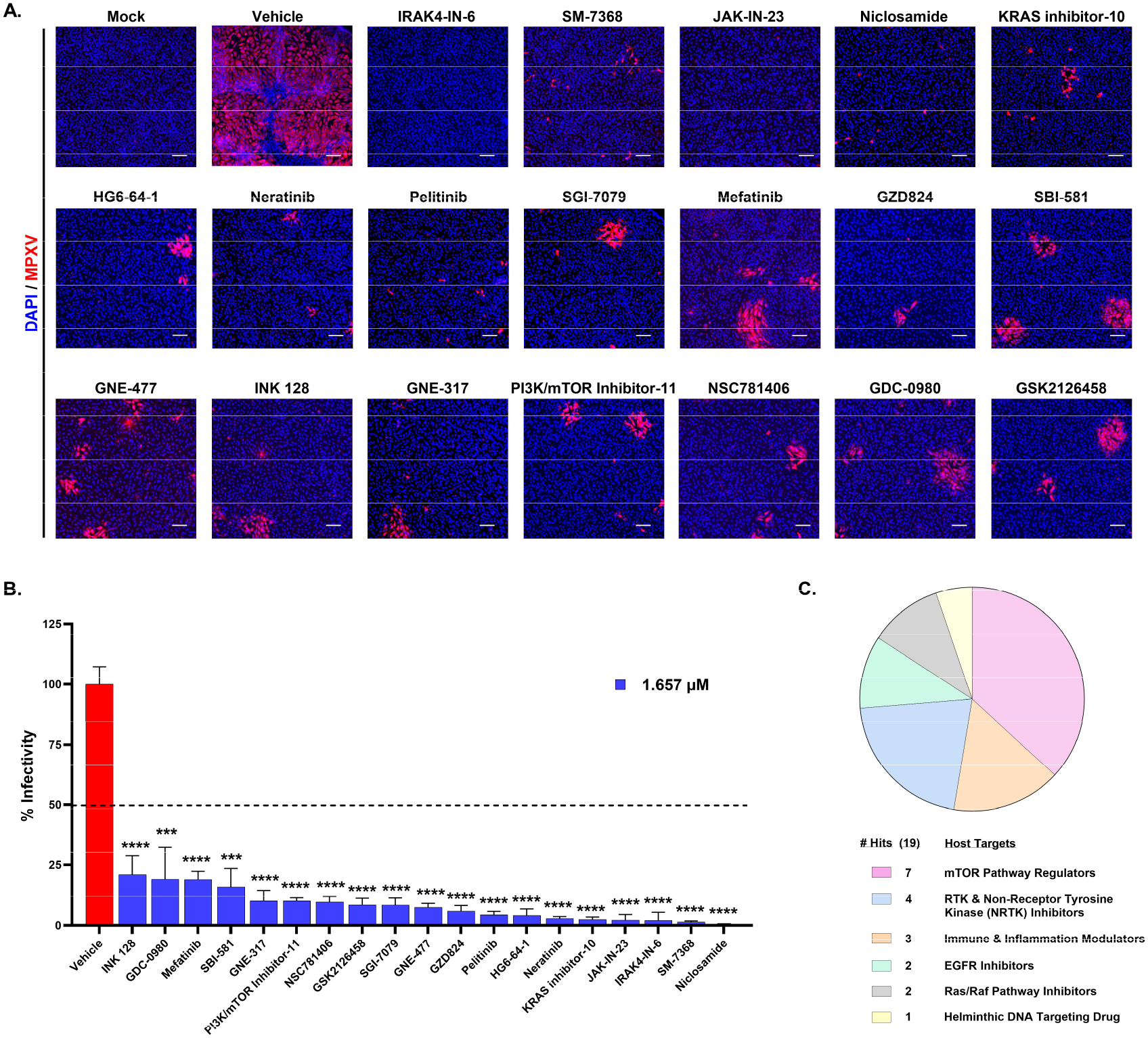
Secondary Screening of MPXV with the Shortlisted Kinase Inhibitor Library from Primary Screen. **(A)** Representative images of ARPE19 cells infected with MPXV at MOI 0.1 and treated with vehicle contrl or antiviral compounds identified in the primary screening for 48 hours post-infection. Blue and red indicate nuclei and MPXV, respectively. Scale bar = 200 µm. **(B)** Quantitative analysis of percent viral infectivity in cells treated with the indicated compounds. All compounds exhibited statistically significant (p < 0.0001) viral inhibition compared to vehicle treatment. The horizontal dotted line represents the 50% infectivity. **(C)** Pie chart illustrating the distribution of host targets associated with the selected drug molecules identified in the secondary screen. The chart represents the allocation of these drug hits across key target pathways based on the connectivity mapping. Quantitative data are presented as mean ± standard deviation (SD). Statistical analysis was performed using ANOVA, followed by Tukey’s post hoc test (*, P < 0.05; **, P < 0.01; ***, P < 0.001; ****, P < 0.0001).

As MPXV has been known to spread sexually and infect the vaginal mucosa, the antiviral activity of the 19 selected compounds was further evaluated in VK2/E6E7 cells – an immortalized human vaginal epithelial cell line^34^. To gain additional insight into toxicity and efficacy across a range of concentrations, a dose-response study was performed using seven concentrations (1.657, 0.828, 0.414, 0.207, 0.103, 0.0517, and 0 µM) in triplicate by pre-treating VK2 cells for 16 hours (Fig. S4). This independent validation identified seven lead drug candidates: SM-7368, IRAK4-IN-6, KRAS inhibitor-10, Niclosamide, GNE-317, INK-128, and PI3K/mTOR inhibitor-11. The dose-response curves showed the half-maximal inhibitory concentration (IC_50_) in the range of 12-50 nM (Fig. 3B). Among the seven, IRAK4-IN-6, SM-7368, and KRAS inhibitor-10 were nontoxic up to a concentration of 1000nM. In contrast, Niclosamide, GNE-317, INK-128, and PI3K/mTOR inhibitor-11 showed toxicity at approximately 80-100nM in VK2 cells (Figure S2). The remaining twelve compounds showed severe toxicity with a very narrow dose range for anti-MPXV activity in this cell type (Fig. S3). Based on these findings, we decided to use the three non-toxic compounds IRAK4-IN-6, SM-7368, and KRAS inhibitor-10, as well as Niclosamide, which showed the strongest viral inhibition (Fig. 2B).

**Figure 3.**
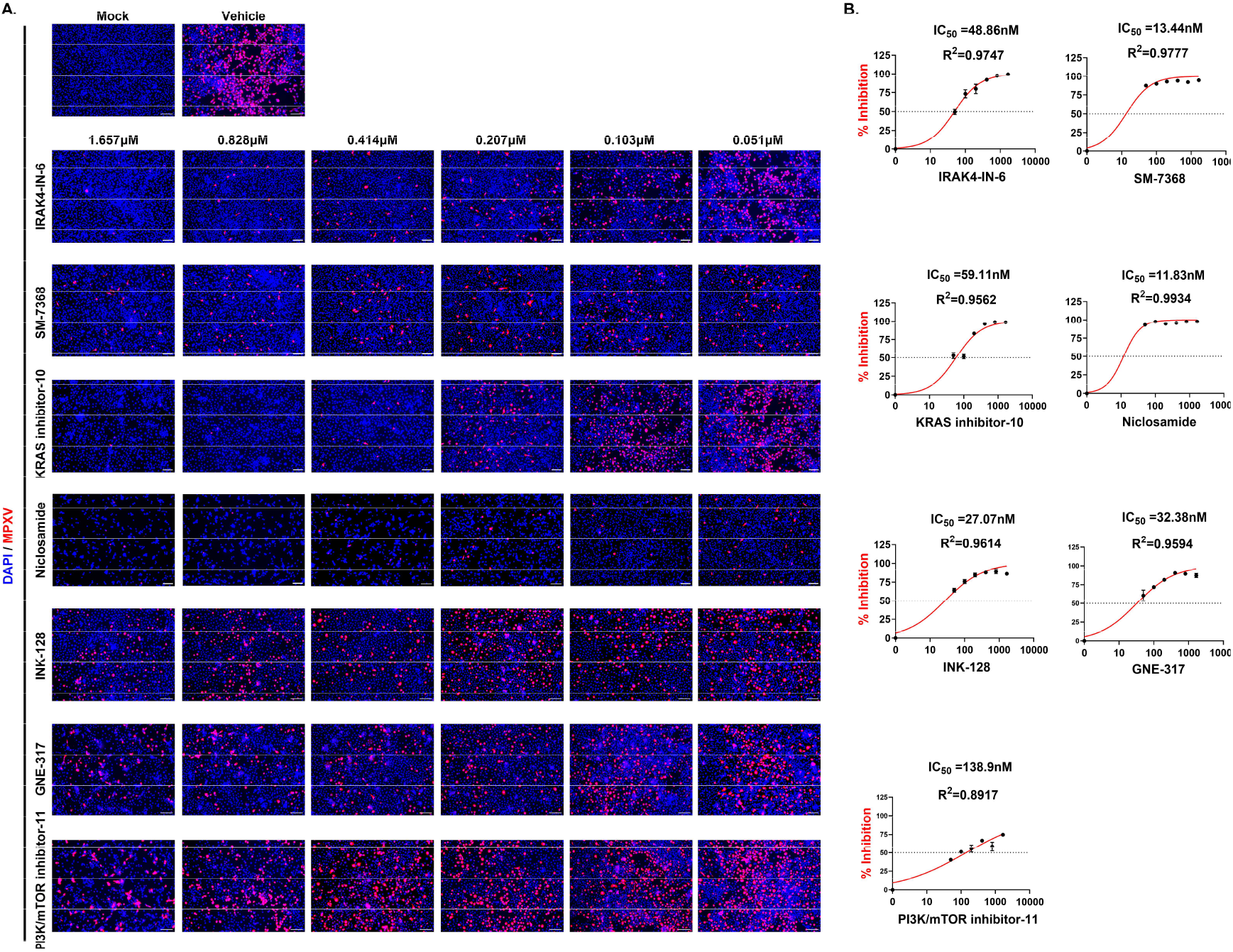
Dose-Response Study of Antiviral Drug Candidates Against MPXV Infection. **(A)** Representative immunofluorescence images of MPXV-infected VK2/E6E7 cells treated with seven selected drug compounds at varying concentrations (1.657, 0.828, 0.414, 0.207, 0.103, and 0.051 µM) for 48 hours post-infection (hpi). Blue and red indicate nuclei and MPXV-A4L antigen, respectively. Scale bar = 100 µm. **(B)** The dose-response curves of the indicated compounds are shown. The percentage of MPXV-Dose–response curves showing percent inhibition of MPXV infection in VK2/E6E7 cells treated with the indicated compounds. Cells were infected with MPXV and fixed at 48 hours post-infection (hpi) for immunofluorescence analysis. Infection levels were calculated as the ratio of MPXV-positive cells (red) to total DAPI-stained nuclei (blue), normalized to vehicle-treated controls. Percent inhibition values were derived as 1 minus the normalized infectivity. Nonlinear regression was used to calculate IC_50_ values, which are displayed along with curve fit R^2^ values. The dotted horizontal line indicates 50% inhibition. Data represent the mean ± standard deviation from triplicate wells.

### Validation of Antiviral Compounds in Primary Human Keratinocytes

The major clinical manifestations of MPXV involve skin lesions similar to those caused by poxvirus, indicating its ability to infect keratinocytes^35^. Thus, we assessed the antiviral activity of the four selected drug candidates (IRAK4-IN-6, SM-7368, KRAS inhibitor-10, and Niclosamide) in primary human epidermal keratinocytes. Cells were pretreated with each drug at 1.0µM for 16 hours, followed by infection with MPXV (MOI 0.1) and immunostaining for MPXV, with cell lysates collected for immunoblotting. Our data showed that all four compounds inhibited viral replication, as evidenced by reduced staining for MPXV antigen (Fig. 4A). Moreover, we observed that all drugs reduced MPXV-induced apoptosis assessed by immunostaining for cleaved caspase-3 (Fig. S5A). Cell viability in the drug-treated groups was above 80% (Fig. S5B). As the compounds IRAK4-IN-6 and SM-7368 have been implicated in regulating inflammatory response^36,37^, we performed Western blot analysis of NF-κB and STING signaling in cell lysates from cells treated with IRAK4-IN-6 and SM-7368 in the presence or absence of MPXV infection (Fig. 4B). Consistent with the reduced MPXV antigen staining (Fig. 4A), viral protein was substantially reduced in cell lysates from the IRAK4-IN-6- and SM-7368-treated groups (Fig. 4B). In addition, our data showed that IRAK4-IN-6 significantly increased phospho-NF-κB and phospho-STING expression, suggesting their immunomodulatory roles during MPXV infection (Fig. 4B).

**Figure 4.**
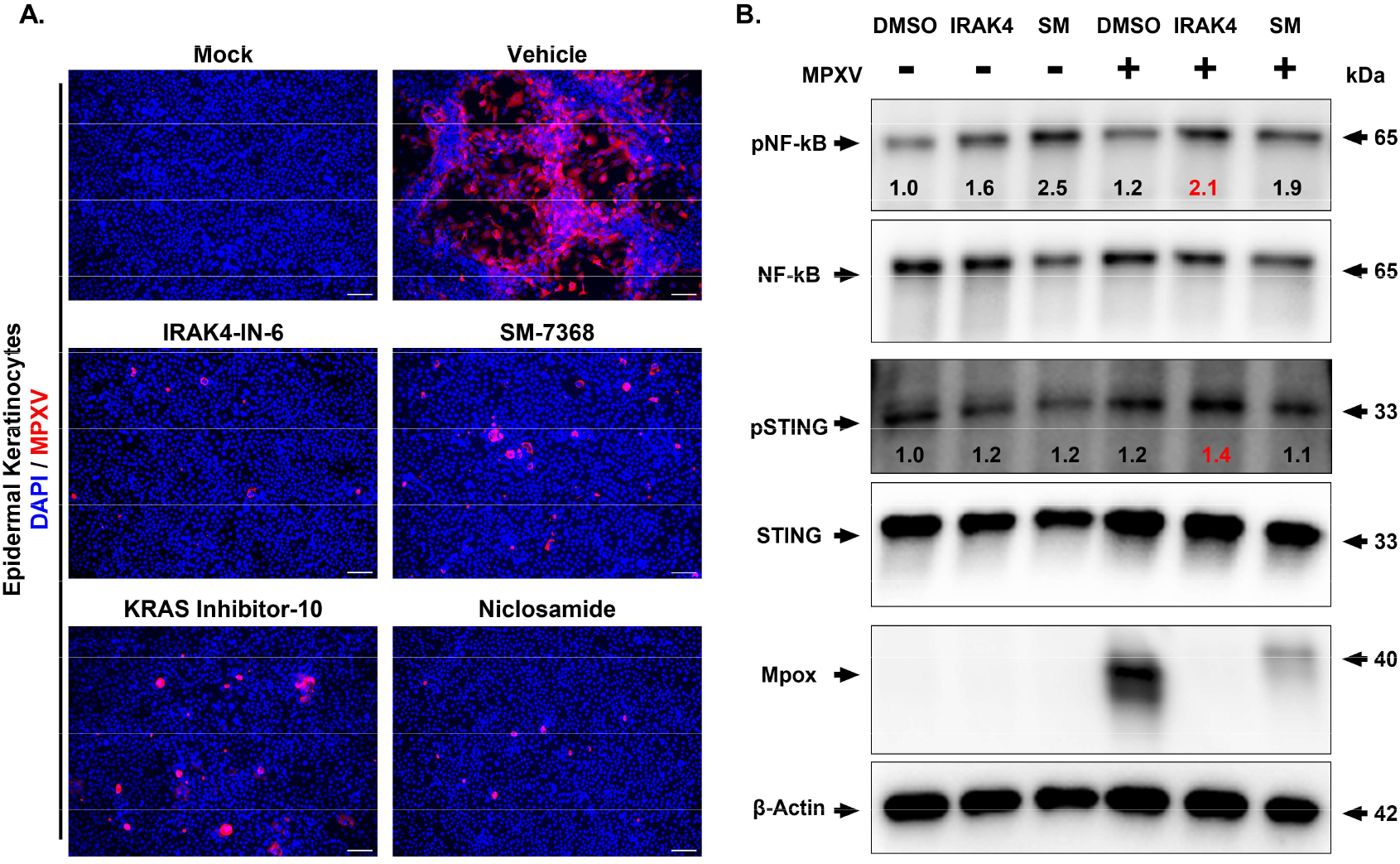
Validation of Selected Drug Compounds Against MPXV in Primary Human Keratinocytes. **(A)** Immunofluorescence analysis of MPXV-infected human epidermal keratinocytes treated with the indicated drug compounds (IRAK4-IN-6, SM-7368, KRAS inhibitor-10, or Niclosamide) at 1µM for 48 hours post-infection (hpi). Blue and red indicate nuclei and MPXV-A4L antigen, respectively. Scale bar = 100 µm. **(B)** Western blot analysis of MPXV-infected cells comparing the effect of drug treatment on MPXV replication, as well as the activation of NF-κB and STING pathways. The normalized band density values of p-STING/STING and p-NFkB/NFkB for each condition are provided.

### *In Vivo* Evaluation of Drug Candidates

Our *in vitro* studies indicated the potent antiviral activity of IRAK4-IN-6, SM-7368, and KRAS inhibitor-10 against MPXV. Next, we developed a dermal mouse model of Mpox infection to test their efficacy *in vivo*. MPXV infection was established via intradermal injection of MPXV (1×10^5^ TCID_50_/mouse) in C57BL/6 mice. All drugs were applied topically twice daily for 7 days, starting eight hours after virus injection. On Days 3 and 12 post-infection, the mice were sacrificed for collecting skin tissues (Fig. 5A). Skin pathology was evaluated by histopathological analysis with hematoxylin and eosin (H&E) staining, which showed that MPXV caused extensive skin lesions at both timepoints, with raised and inflamed edges around the infection site along with increased infiltration of inflammatory cells, such as neutrophils (Fig. 5B and S6). However, none of the mice in the IRAK4-IN-6-treated group had visible skin lesions or neutrophil infiltration at Day 3 or Day 12 post-infection (Fig. 5B and S6). In the case of SM-7368 and KRAS inhibitor-10 treatment, mice exhibited some low-grade inflammation and lesions at Day 3, but these were fully resolved by Day 12. Immunostaining of skin tissue showed reduced levels of MPXV antigen in the drug-treated groups compared to vehicle-treated skin tissue, both at Day 3 and Day 12 (Fig. 5C, 5D, and S6). Together, these results demonstrate that IRAK4-IN-6, SM-7368, and KRAS inhibitor-10 exhibit potent antiviral activity against MPXV and reduce viral-induced pathology in skin tissue, making them promising candidates for further pre-clinical drug development.

**Figure 5.**
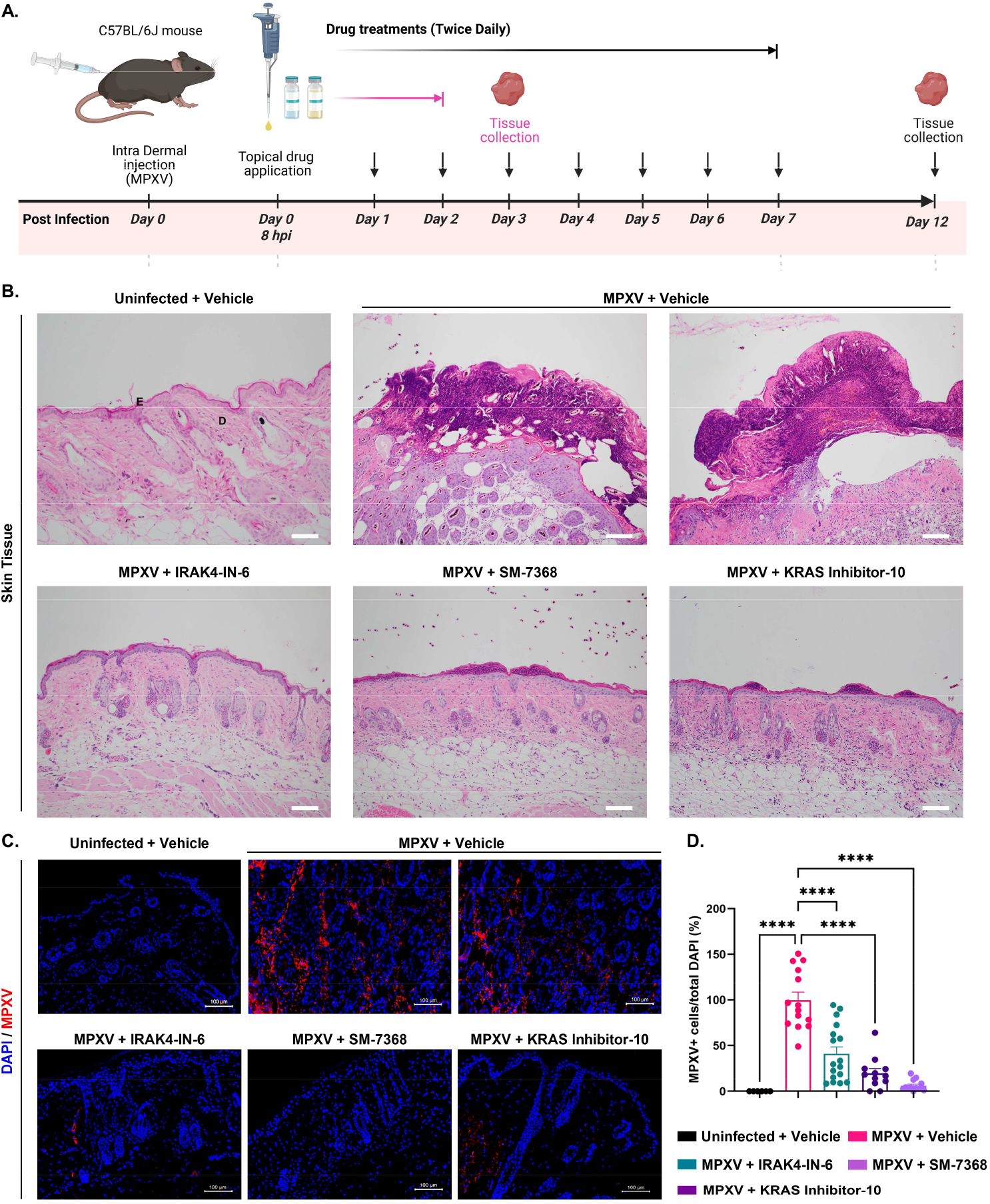
Testing the Efficacy of MPXV Antiviral Compounds in a Dermal Mpox Mouse Model. **(A)** Overview of the experimental timeline showing intradermal MPXV infection (Day 0), topical drug application period, and skin tissue harvest on Day 3, the focus of this main figure, highlighted in pink. **(B)** Histological examination of skin sections collected on Day 3 from MPXV-infected mice treated with topical formulations of IRAK4-IN-6, SM-7368, or KRAS inhibitor-10. Hematoxylin and eosin (H&E) staining was performed, and images were captured at 10× magnification. Labels “E” and “D” denote the epidermis and dermis, respectively, for reference. **(C)** Immunohistochemistry analysis of MPXV-infected mouse skin collected on Day 3 post-infection. Sections were stained for MPXV A4L viral antigen (red) and counterstained with DAPI (blue) to visualize nuclei. **(D)** Quantification of MPXV-positive cells expressed as percent infectivity. Statistical significance was assessed using one-way ANOVA followed by Tukey’s multiple comparisons test (*P < 0.05; **P < 0.01; ***P < 0.001; ****P < 0.0001).

## DISCUSSION

With the cessation of smallpox vaccination decades ago, the proportion of people with cross-immunity for MPXV has rapidly declined. In addition, increased international travel, a warming climate, and population growth present the conditions for increased global virus spread and mutations among MPXV strains. With the emergence of novel strains of MPXV and increased human-to-human spread globally, there is an urgent and unmet need for MPXV-specific antiviral agents.

To rapidly establish new effective drugs against MPXV, we decided to evaluate small molecules targeting host cell pathways, utilizing a carefully curated chemical library for screening to shorten the generally costly and lengthy process of developing direct-acting antivirals. Targeting the host cellular pathways essential for virus propagation rather than the virus itself may also reduce antimicrobial resistance. Once identified, compounds that are already safety-tested in humans can be rapidly deployed for clinical use.

Potential direct-acting antiviral drugs mostly target viral replication by interfering with their genomic DNA or RNA synthesis, as well as inhibiting viral protein function. Nucleoside analogs such as cidofovir exhibit a mode of action of competing with the available pool of cellular nucleotides, thus blocking viral replication *in vitro* and in animal models^38,39^. Following the Mpox outbreak in 2022, cidofovir was tested in clinical trials but had too many serious side effects, such as kidney toxicity^40,41^. Brincidofovir, a cidofovir derivative, had potent antiviral efficacy in cell culture and mice, but it induced similarly severe adverse effects^42,43^. Ribavirin, another nucleoside analog with broad-spectrum antiviral efficacy, can block MPXV replication *in vitro* and in mouse models^44^. However, ribavirin is known to cause severe bone marrow suppression in humans. Moreover, tecovirimat (TPOXX) is used off-label to treat topical Mpox lesions. However, the recent STOMP trial showed it to be ineffective, and another study showed cases of tecovirimat resistance in severely immunocompromised Mpox patients with prior tecovirimat use^31,33^.

In this study, we utilized relevant *in vitro* human epithelial cell model systems to identify potential therapeutic compounds targeting MPXV and verified their therapeutic potential in an *in vivo* dermal Mpox mouse model. Our initial results yielded compounds inhibiting the EGFR, PI3K-mTOR, Ras/Raf, and NF-κB pathways. We found that IRAK4-IN-6, SM-7368, and KRAS inhibitor-10 exhibited significant antiviral activity against MPXV and ameliorated Mpox-related skin lesions in mice.

IRAK4-IN-6 is a potent inhibitor of IRAK4, a key player in Toll-like receptor (TLR) and IL-1R innate immune signaling^45–47^. TLR2 and TLR9 have been involved in eliciting innate immune responses to MPXV^48,49^, suggesting that TLR signaling can be targeted for Mpox treatment. Our data showed that IRAK4-IN-6 activated NF-κB and STING pathways by phosphorylating these two proteins, likely potentiating an antiviral response. SM-7368 is an inhibitor of NF-κB, targeting downstream of MAPK p38 activation, which has also been shown to inhibit IL-1β-mediated induction of SDC4 promoter activity^50^. SM-7368 has also been found to inhibit TNF-α-induced matrix metalloproteinase-9 (MMP-9)^51^, which, as such, has led to investigations into its potential use as a chemotherapeutic agent against tumor invasion and metastasis^52^. KRAS inhibitors have commonly been investigated for their potential role as chemotherapeutic agents, primarily against colorectal and lung cancers, due to their ability to promote Type I IFN and STING pathway expression^53,54^. IFN-β has been shown to significantly reduce Clade I MPXV infection *in vitro*^55^, suggesting the potential involvement of Type I IFN pathways as an important host-directed antiviral target. Considering the possible roles of these drugs in modulating innate immune function, further investigations are warranted to establish their mode of action in inhibiting viral proliferation.

The role of EGFR inhibitors is primarily to target host kinase receptors^56^. However, considering that MPXV encodes D3R, an EGF analog^22,56^, these compounds may act as competitive inhibitors of viral EGF activation of the host EGFR signaling pathway. Interestingly, the EGFR inhibitor gefitinib was found to reduce vaccinia and cowpox virus replication in cell culture^57^. Niclosamide is an FDA-approved orally administered compound that is widely used clinically for the treatment of helminthic infection. However, the compound can inhibit the Wnt/β-catenin and STAT3 pathways mediating ovarian cell proliferation^58^, as well as the synthesis of cellular DNA and change the cellular filamentous actin cytoskeleton network^59^, thus limiting its utility in systemic infections. However, it has the potential to be further developed into a topical treatment for Mpox skin lesions.

Beyond established tropism for skin epithelial cells^60–62^, MPXV has been shown to infect cells of the eye^7^, respiratory tract^60–62^, central nervous system^63^, and immune system^22,64^. Considering the irritation due to Mpox skin lesions and their role in promoting MPXV transmission^65^, there is a need to develop effective intradermal and topical treatments for Mpox. As such, our investigation focused on an intradermal route of delivery. However, with recent evidence of newer MPXV strains capable of sexual transmission^4^/11/2025 10:08:00 PM, with many cases showing the presence of anogenital Mpox lesions^66^, there is a need to better elucidate the tropism of MPXV for cells of the anogenital epithelium, as well as directing treatments specifically for the anogenital region. We observed potent infection by MPXV of human vaginal epithelial cells, as well as the efficacy of these drug compounds in mitigating this infection *in vitro*. As such, we plan to further evaluate these compounds for treating systemic Mpox via oral or parenteral routes of administration.

This is the first study to evaluate a comprehensive kinase inhibitor drug library to evaluate potential therapeutic agents specifically against MPXV. It should be noted that this study only evaluated the efficacy of these compounds against Clade II MPXV strains, and further investigations are needed to assess their ability to treat Clade Ia and newer Clade Ib strains. Given the 96% genetic similarity between the smallpox virus and MPXV, some of our compounds with antiviral activity may also be effective against this dangerous pathogen. These compounds represent valuable therapeutic candidates to retain in preparation for potential re-emergent or newly emergent poxviral outbreaks. Future studies should focus on evaluating combination therapies involving two or more drugs to develop the most potent anti-Mpox treatment cocktails for various virus-related indications. Taken together, our study using high-throughput drug screening represents a promising step toward identifying effective treatments for Mpox. This is especially important given the recently reported limitations of tecovirimat and the ongoing need for targeted antivirals with improved safety profiles and a lower risk of resistance. Such treatments are critical for addressing this highly infectious and rapidly evolving virus.

## Acknowledgments

We are grateful to Barbara Dillon, UCLA High Containment Program Director, for BSL-3 work. The following reagent was obtained through BEI Resources, NIAID, NIH: Monkeypox Virus, hMPXV/USA/MA001/2022 (Lineage B.1, Clade IIb), NR-58622. This study was partially funded by NIH grants 1R01EY032149-01, 5R01AI163216-02, and 1R01DK132735-01 to VA, and R01EY026964, R01EY027381, and R01EY035499 to AK.

The UCLA Molecular Screening Shared Resource is supported by the Jonsson Comprehensive Cancer Center and award number P30CA016042 by the National Cancer Institute (NIH).

## Author contributions

AVJ, AZ: Collection and/or assembly of data. Data analysis and interpretation, and Manuscript writing.

NC, SS, BN, RP, NS, EG, AJ, JT, LF, NL, CT, SJ, MP, SWF, AD, MB, GG: Experimental design, Conducted experiments, Data analysis, interpretation, and Manuscript writing.

AK, RD, VA: Conception and design, Data analysis and interpretation, Manuscript writing, and Final approval of manuscript.

## Competing interests

The authors declare no competing interests.

## Data and materials availability

All mentioned and relevant data regarding this study is available from the above-listed authors. In addition, supplementary information is available for this paper. Correspondence and requests for data and materials should be addressed to lead contact, Vaithilingaraja Arumugaswami.

## METHODS AND MATERIALS

### Ethics Statement

This study was performed in strict accordance with the recommendations of UCLA. All MPXV live virus experiments were performed at the UCLA BSL-3 High Containment facility. The experiments were approved by the UCLA Institutional Biosafety Committee (IBC) and the UCLA Institutional Animal Care and Use Committee (IACUC).

### Cells

Human retinal pigmented epithelial cell line (ARPE-19) (ATCC, CRL-2302) was cultured in DMEM: F-12 Medium, supplemented with 10 % fetal bovine serum (FBS),1 % MEM Non-Essential Amino Acids Solution (MEM NEAA), 1 % L-glutamine (L-glu) and 1 % penicillin/streptomycin (P/S). The VK2/E6E7 (VK2) human vaginal epithelial cell line (ATCC, CRL-2616) was maintained in keratinocyte serum-free medium (KSFM) supplemented with 0.1 ng/mL human recombinant epidermal growth factor (EGF), 0.05 mg/mL bovine pituitary extract (BPE), and 100 U/mL penicillin-streptomycin. The NHEK-Neo cells (Neonatal Human Epidermal Keratinocytes) (Lonza, #00192906) were cultured in KGM Gold Keratinocyte Growth Medium BulletKit, consisting of KBM Gold Basal Medium supplemented with KGMTM Gold SingleQuots, as per the manufacturer’s instructions. All cells were maintained at 37°C in a humidified incubator with 5% CO_2_.

### Viruses

MPXV virus hMPXV/USA/MA001/2022 (Lineage B.1, Clade IIb) stock was received from BEI Resources. We have passaged the virus once in Vero E6 cells. Then, sequence-verified viral stocks were aliquoted and stored at −80°C. The virus titer was measured in Vero E6 cells with the established TCID_50_ assay.

### Viral Titer by TCID50 (Median Tissue Culture Infectious Dose) *assay*

Viral production by infected cells was quantified by the TCID50 assay, as previously described with modifications^48^. Vero E6 cells (density of 5×10^3^ cells/well) were plated in 96-well plates. The next day, viral culture media were serially diluted 10-fold (10^1^ to 10^8^) and added to Vero E6 cells. These cells were incubated at 37ºC with 5% CO_2_. 3 to 4 days post-infection, the cells were examined for viral CPE. The wells positive for viral infection were identified for each dilution. Then, the dilutions immediately above and below 50% of viral inhibition were determined. TCID50 was calculated based on the method of Reed and Muench.

### Viral Infection

Cells were plated at 1×10^4^ or 3×10^3^ cells per well using 96-well or 384-well plates, respectively. 25 µl viral inoculum of MPXV was added onto the cells at a multiplicity of infection (MOI) of 0.1 using serum-free base media. Infected cells were incubated at 37ºC with 5% CO_2_. Cells were then fixed at selected time points with 4% paraformaldehyde (PFA). As needed, samples were then collected, and RIPA buffer was used for protein analysis.

### Drug Compounds, Infections, and Screening

The kinase inhibitor library, composed of 2750 small molecules inhibiting host kinases, is curated at the UCLA Molecular Screening Shared Resource. The compounds were also obtained from MedChemExpress and Selleckchem. All compounds were provided as lyophilized and were then reconstituted in DMSO. Compounds were then aliquoted and stored at either −80°C or room temperature in dry conditions. For all drug screening studies, the indicated drugs were co-seeded with cells and incubated for 24 hours prior to MPXV infection (MOI 0.1). The final concentration of DMSO was maintained at 0.5% in all wells, regardless of the compound concentration. The cells were fixed with 4% PFA at 48hpi for downstream immunofluorescence (IFA) analysis. The plates were transferred to an Image Xpress confocal microscope and imaged (20x, Nikon ELWD Objective, SFluor, NA 0.45, 2-4 sites per well or similar). Infected cells were quantified using the Molecular Devices Cell Scoring Module, typically using 1500 grey scales as cutoff of 5-20µm for nuclei detection and at least 1000 grey scales as cutoff and a size of 15-40µm for the detection of viral proteins within the cytosol. The total number of nuclei and percentage of infected cells were exported, as well as averages for data mining. Hits were detected using a 3 z-score cutoff and subjected to further analysis.

### Animal Study

Six-to eight-week-old female C57BL/6 mice were used for this study. The animals were maintained and housed in the UCLA Vivarium under BSL-3 precautions. Prior to infection, the hair on the top-lower back region (above flank) was gently removed using a chemical depilatory method. Each mouse received subdermal inoculation of MPXV (1×10^5^ TCID_50_/animal) in 20µL volume in four individual neighboring sites in the shaved region. At 8hpi, the animals received their first dose of vehicle or drug (5mg/kg) treatment via topical application (20µL) on the infected site using a pipette. The vehicle (DMSO) and the drug stocks were diluted in saline for skin application. The animals received drug treatment twice daily until Day 7 post-infection. The animals were monitored for changes in body weight and health conditions daily. The animals were euthanized on Day 3 and Day 12 endpoints. The skin tissues were fixed in 4% paraformaldehyde for histological analysis.

### Immunofluorescence Assay

Cells were fixed with 4% PFA or 80% methanol for 20 minutes at room temperature or −20°C, respectively. The cells were washed three times with 1X PBS and permeabilized by incubating in blocking buffer (0.3% Triton X-100, 2% BSA, 5% Goat Serum, 5% Donkey Serum in 1 X PBS) for 1 hour at room temperature. For immunostaining, cells were incubated overnight at 4ºC with each primary antibody, then washed with 1X PBS three times and incubated with respective secondary antibody for 1 hour at room temperature. The cell nuclei were stained with DAPI (4’,6-Diamidino-2-Phenylindole, Dihydrochloride) (Life Technologies) at a dilution of 1:5000 in 1X PBS. Image acquisition was done using Leica DM IRB fluorescent microscopes (Leica Microsystems, Illinois, USA).

### Immunohistochemistry

The skin tissue samples were fixed with 4% PFA, paraffin-embedded, and cut into 5-μm sections, and indirect immunofluorescence (IFA) staining was performed for the detection of the MPXV antigen. The sections were deparaffinized and rehydrated, followed by 15 minutes heat-induced antigen retrieval with Citrate EDTA buffer pH 8.0. Next, the sections were permeabilized and blocked using 10% normal goat serum with 0.5% Triton X-100 for an hour at room temperature in a humidified chamber, followed by overnight incubation at 4°C with mouse Anti-Vaccinia Virus E3L monoclonal antibody (1:100). The sections were washed thrice in 1X PBS and incubated with Donkey anti-Mouse IgG Secondary Antibody, Alexa Fluor 594 (1:200; ThermoFisher Scientific) for an hour at 37C. The skin sections were extensively washed with PBS (4 washes, 5 min each), and the slides were mounted in Vectashield antifade mounting medium (Vector Laboratories, Burlingame, CA). The images were acquired using a Keyence microscope (Keyence, Itasca, IL).

### Cell Viability Assay

Cell viability and intracellular ATP levels were assessed using the CellTiter-Glo Luminescent Cell Viability Assay (Promega), following the manufacturer’s instructions. VK2/E6E7 cells (100 µL per well) were seeded directly into 96-well flat-bottom plates along with six different concentrations of the selected compounds (SM-7368, IRAK4-IN-6, KRAS inhibitor-10, Niclosamide, GNE-317, INK-128, and PI3K/mTOR inhibitor-11). Each drug concentration was plated in quadruplicate wells at the time of seeding. After 48 hours of incubation, 100 µL of CellTiter-Glo reagent was added to each well and incubated for 10 minutes at room temperature. The mixture was gently pipetted to ensure proper mixing, and 100 µL of the cell–reagent solution was transferred to a white-bottom 96-well plate for detection. Luminescence was measured using a GloMax® Discover plate reader (Promega). Percent cell viability was calculated relative to wells treated with DMSO alone.

### Image Analysis/Quantification

Images of tissue culture cells were obtained using the Leica DM IL LED Fluo and Leica LAS X Software Program. Four to six images were captured per well at 48hpi for each condition. These images were quantified using ImageJ’s plugin (Multipoint and Cell Counter). The positively stained cells were counted by a double-blinded approach.

### Statistics and Data Analysis

GraphPad Prism, version 10, was used for graph generation and statistical analysis. Data was then analyzed for statistical significance using an unpaired student’s *t*-test to compare two groups (uninfected vs. infected) or a non-parametric t-test (Mann-Whitney Test). All data is representative of two or more experiments with three to four biological replicates. All statistical testing was performed at two-sided alpha levels of 0.05, 0.01, or 0.001.

## Notes

### Competing Interest Statement

The authors have declared no competing interest.

