## Supplementary Figures for "Drug screen reveals new potent host-targeted antivirals against Mpox virus"

#### SUPPLEMENTAL FIGURES AND LEGENDS

##### Supplementary Figure 1

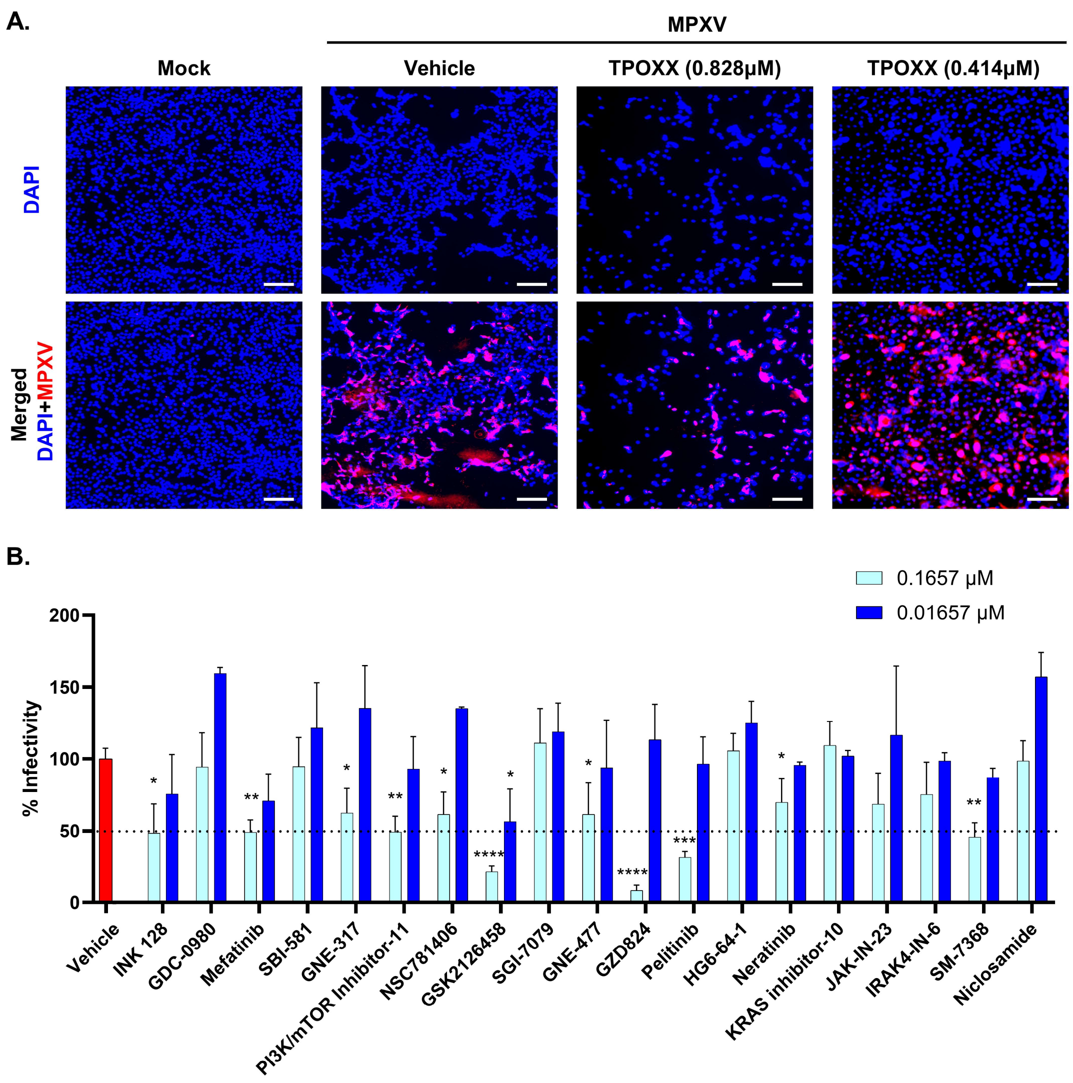

***Supplementary Figure 1. A)*** *Immunofluorescence images of MPXV-infected retinal pigment epithelial cells, ARPE-19 (red) treated with Tecovirimat (TPOXX) at 0.828 μM and 0.414 μM for 48 hours post-infection (hpi). DAPI (blue) was used to counterstain nuclei. Scale bar = 100 μm.* ***B)*** *Quantification of MPXV-infected cells based on immunofluorescence imaging following treatment with 19 selected compounds from the secondary screening at 0.1657 μM (light blue) and 0.01657 μM (dark blue) for 48 hours post-infection (hpi). Percent infectivity was calculated as the ratio of MPXV-positive cells (red) to total DAPI-stained nuclei (blue) and normalized to the average infectivity of the vehicle-treated control group. The dashed horizontal line indicates the 50% infectivity threshold. Data represent the mean ± standard deviation (SD) from image-based quantification across three wells per condition. Statistical analysis was performed using one-way ANOVA, followed by Tukey's post hoc test (*P < 0.05; **P < 0.01; ***P < 0.001; ***P < 0.0001).*

#### Supplementary Figure 2

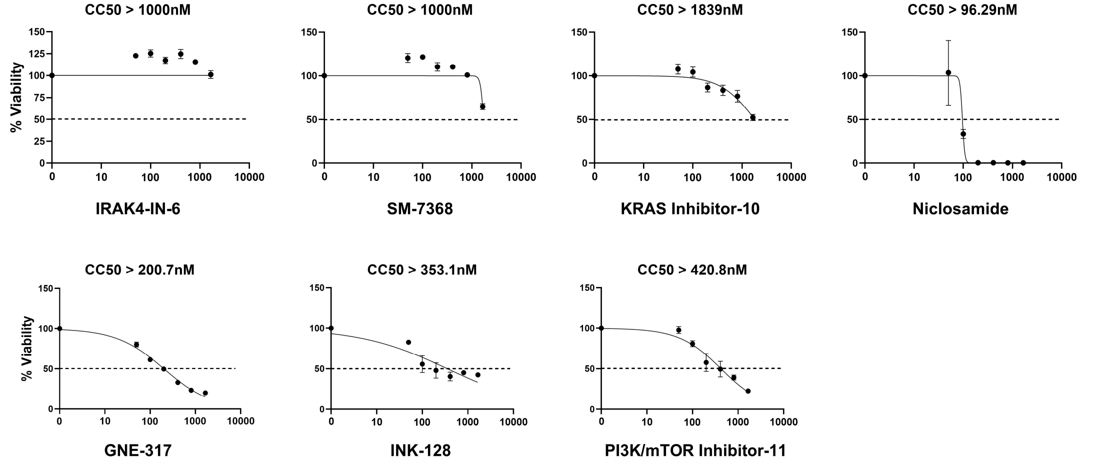

***Supplementary Figure 2.*** *Cell viability of VK2/E6E7 human vaginal epithelial cells treated with seven selected drug compounds was assessed using the CellTiter-Glo Luminescent Cell Viability Assay. Cells were incubated with each compound for 48 hours, and luminescence was measured to quantify intracellular ATP levels, which correlate with metabolically active (viable) cells. Dose–response curves were generated to determine the half-maximal cytotoxic concentration ($CC_{50}$) for each compound. IRAK4-IN-6, SM-7368, and KRAS inhibitor-10 exhibited low cytotoxicity with $CC_{50}$ values >1000 nM, while Niclosamide demonstrated higher cytotoxicity ($CC_{50}$ = 96.29 nM). GNE-317, INK-128, and PI3K/mTOR Inhibitor-11 showed moderate cytotoxic effects with $CC_{50}$ values of 200.7 nM, 353.1 nM, and 420.8 nM, respectively. The dotted horizontal line indicates the 50% viability threshold, and error bars represent the standard deviation (SD).*

### Supplementary Figure 3

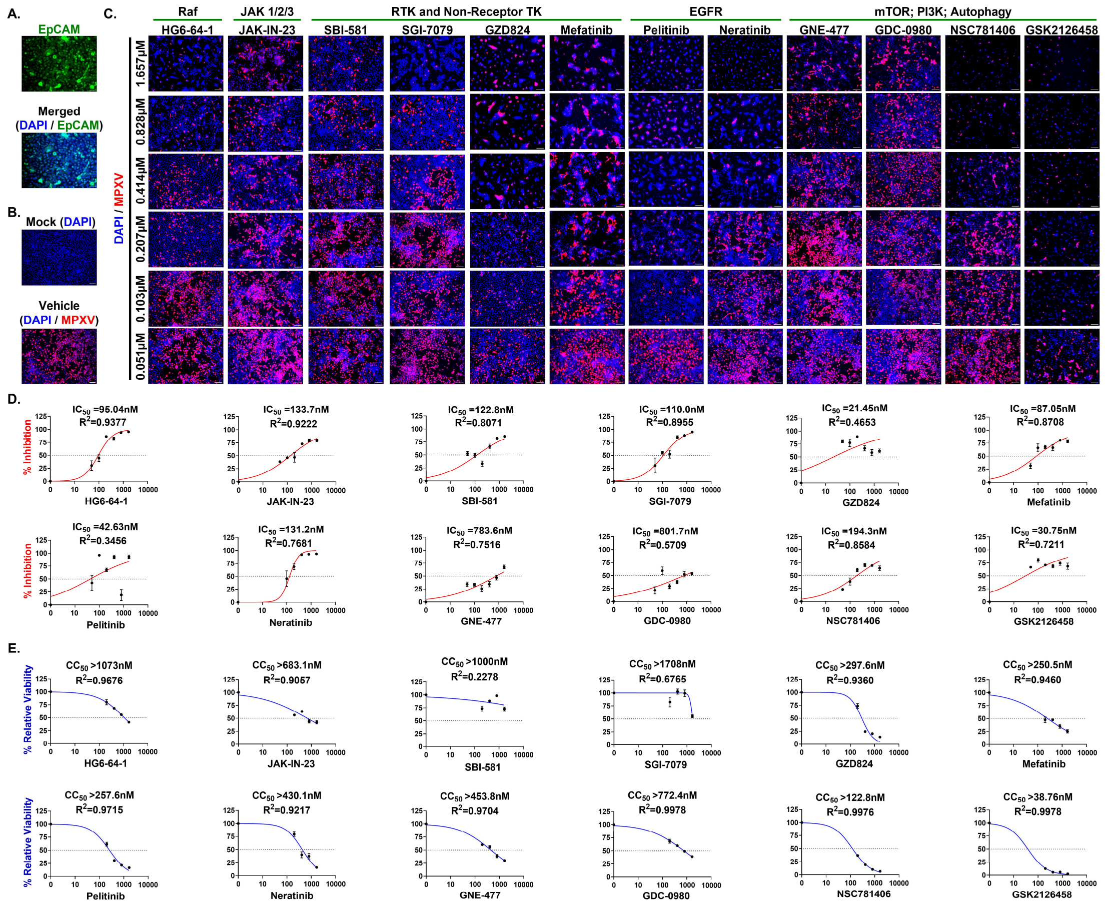

***Supplementary Figure 3. A)*** *Immunofluorescence staining of the epithelial cell marker EpCAM (green) in human vaginal epithelial (VK2/E6E7) cells. Nuclei were counterstained with DAPI (blue). The merged image confirms epithelial cell identity. Scale bar = 100 µm.* ***(B)*** *Control conditions: Mock (uninfected) and Vehicle (MPXV-infected, untreated) wells stained for MPXV A4L viral antigen (red) and DAPI (blue) to visualize nuclei. These controls serve as reference for infection and treatment comparison. Scale bar = 100 µm.* ***(C)*** *Immunofluorescence images of MPXV-infected VK2/E6E7 cells following treatment with 12 drug compounds at six dose concentrations (1.657, 0.828, 0.414, 0.207, 0.103, and 0.051 µM). Cells were fixed at 48 hours post-infection (hpi) and stained for MPXV A4L antigen (red) and nuclei (DAPI, blue). The compounds are grouped by host target classes: Raf, JAK1/2/3, RTK and Non-Receptor Tyrosine Kinases, EGFR, and mTOR/PI3K/Autophagy, as labeled above each set. Scale bar = 100 µm.* ***(D)*** *Dose–response curves showing percent inhibition of MPXV infection in VK2/E6E7 cells treated with 12 compounds across six concentrations. Cells were infected with MPXV and fixed at 48 hours post-infection (hpi) for immunofluorescence analysis. Infection levels were calculated as the ratio of MPXV-positive (red) to DAPI-stained (blue) nuclei, normalized to vehicle controls. $IC_{50}$ values were determined using nonlinear regression. The dotted line indicates 50% inhibition.* ***(E)*** *Dose–response curves showing relative cell viability for the drug treatments. Viability was assessed based on DAPI-positive nuclei counts, normalized to mock-treated (uninfected) controls. $CC_{50}$ values were calculated using the top four concentrations for curve fitting. Data are presented as mean ± SD from triplicate wells. The dotted line indicates 50% viability.*

#### Supplementary Figure 4

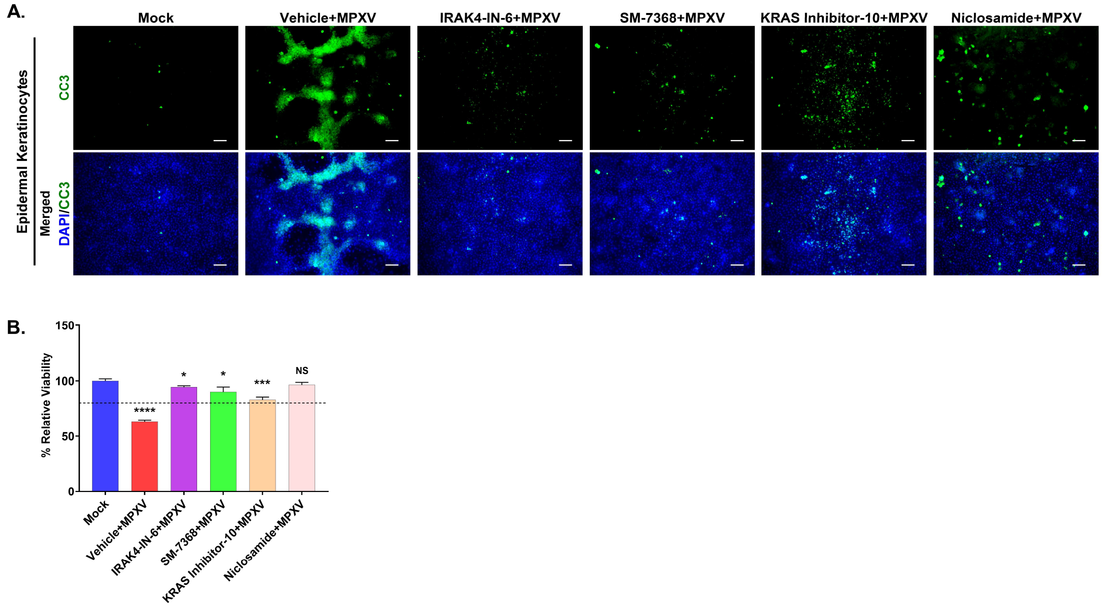

***Supplementary Figure 4. A)*** *Immunofluorescence staining of human epidermal keratinocytes pre-treated with four selected drugs (IRAK4-IN-6, SM-7368, KRAS inhibitor-10, and Niclosamide) at 1.65μM, followed by MPXV infection and incubation for 48 hours post-infection (hpi). Cleaved caspase-3 (CC3, green) was used as a marker for apoptotic cells. Scale bar=100 μm.* ***(B)*** *Quantification of relative cell viability in MPXV-infected keratinocytes treated with four selected drug compounds. Viability was assessed by counting DAPI-stained nuclei in each condition and normalizing to the average DAPI cell count of mock (uninfected, untreated) wells. Data are presented as mean ± standard deviation. Statistical analysis was performed using ANOVA, followed by Tukey's post hoc test (*P < 0.05; **P < 0.01; ***P < 0.001; ****P < 0.0001; NS, not significant).*

#### Supplementary Figure 5

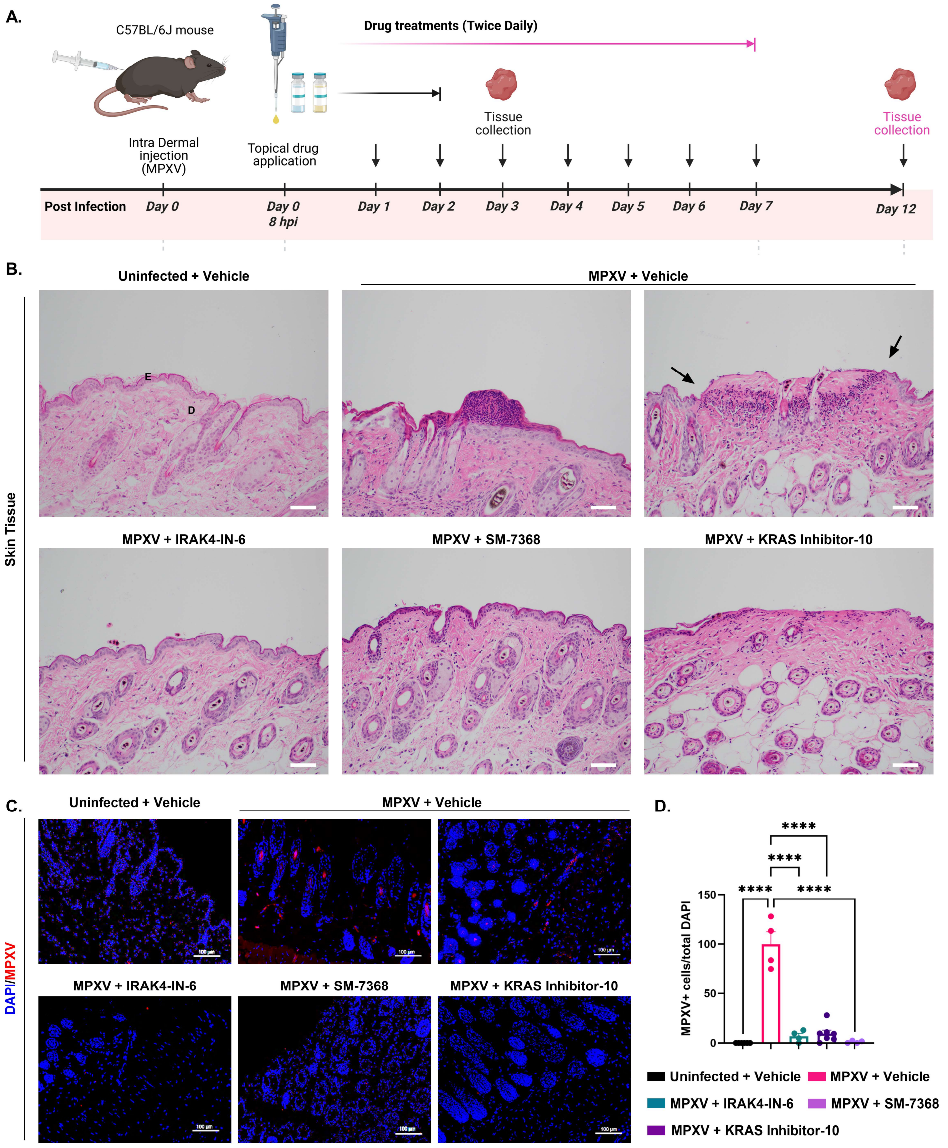

***Supplementary Figure 5. A)*** *Schematic representation of the experimental workflow, illustrating the MPXV infection timeline, drug administration period, and tissue collection timepoints with day 12 highlighted in pink to indicate the focus of this supplementary figure.* ***(B)*** *Histopathological analysis of Day12 collected skin tissues from MPXV-infected mice topically treated with IRAK4-IN-6, SM-7368, and KRAS inhibitor-10. Representative H&E-stained images were captured using a 10X objective lens. MPXV-infected (vehicle-treated) mouse skin showed scab formation, leading to skin erosion (indicated by black arrows), with heavy inflammatory cell infiltrate. Drug treatment resulted in the resolution of these Mpox skin lesions with reduced inflammation. Labels "E" and "D" indicate the epidermis and dermis layers, respectively, as general reference points for skin morphology.* ***(C)*** *Immunohistochemistry staining of MPXV-infected mouse skin sections harvested on Day 12 post-infection. Tissue sections were stained for MPXV (MPXV A4L) (red) and DAPI (blue).* ***(D)*** *Viral antigen-positive cells were quantified, and percent infectivity was calculated for the graph. Statistical analysis was performed using ANOVA, followed by Tukey's post hoc test (*, P < 0.05; **, P < 0.01; ***, P < 0.001; ****, P < 0.0001).*

#### SUPPLEMENTARY TABLES

**Supplementary Table 1.** This table includes all 138 drug compounds identified in the primary screening, along with their target pathways and molecular structures.

**Supplementary Table 2.**

| REAGENT/RESOURCE | SOURCE | IDENTIFIER |
|---|---|---|
| **Antibodies** | | |
| Mpox virus A4L antibody [HL2555] | GeneTex | Cat#GTX638927 |
| Phospho-NF-κB p65 (Ser536) (93H1) Rabbit mAb | Cell Signaling | Cat#3033 |
| NF-κB p65 (D14E12) XP® Rabbit mAb | Cell Signaling | Cat#8242 |
| Phospho-STING (Ser366) (E9A9K) Rabbit mAb | Cell Signaling | Cat#50907 |
| STING (D2P2F) Rabbit mAb | Cell Signaling | Cat#13647 |
| EpCAM (D9S3P) Rabbit mAb #14452 | Cell Signaling | Cat#14452 |
| Cleaved Caspase-3 (Asp175) Antibody #9661 | Cell Signaling | Cat#9661 |
| Goat anti-Mouse IgG (H+L) Cross-Adsorbed Secondary Antibody, Alexa Fluor 555 | Thermo Fisher Scientific | Cat#A-21422 |
| Goat anti-Rabbit IgG (H+L) Cross-Adsorbed Secondary Antibody, Alexa Fluor™ 488 | Thermo Fisher Scientific | Cat#A-11008 |
| Monoclonal Anti-Beta-Actin, Clone AC-74 produced in mouse | MilliporeSigma | Cat#A2228 |
| **Bacterial and Virus Strains** | | |
| hMPXV/USA/MA001/2022 (Lineage B.1, Clade IIb) | BEI Resources | Cat#NR-58622 |
| **Chemicals, Peptides, and Recombinant Proteins** | | |
| Regular Fetal Bovine Serum | Corning | Cat#35010CV |
| Eagle's Minimum Essential Medium (MEM) | Corning | Cat#10009CV |
| Penicillin-Streptomycin (10,000 U/mL) | Gibco | Cat#15140122 |
| L-Glutamine (200 mM) | Gibco | Cat#25030081 |
| MEM Non-Essential Amino Acids Solution(100X) | Gibco | Cat#11140050 |
| KGM™ Gold Keratinocyte Growth Medium BulletKit™ | Lonza | Cat#00192060 |
| DMEM/F-12, HEPES | Thermo Fisher Scientific | Cat#11330032 |
| SM-7368 | MedChemExpress | Cat#HY-116626 |
| IRAK4-IN-6 | MedChemExpress | Cat#HY-130253 |
| KRAS inhibitor-10 | MedChemExpress | Cat#HY-138295 |
| Tecovirimat | Selleckchem | CAT#S3380 |
| Methanol, Optima™ LC/MS Grade, Fisher Chemical | Fisher Scientific | Cat#A456-500 |
| Dimethyl sulfoxide | MilliporeSigma | Cat#D2650 |
| 16% Paraformaldehyde (formaldehyde) aqueous solution | Electron Microscopy Sciences | Cat#15710 |
| Dulbecco's Phosphate-Buffered Salt Solution 1X | Corning | Cat#21030CV |
| DAPI (4',6-Diamidino-2-Phenylindole, Dihydrochloride) | Thermo Fisher Scientific | Cat#D1306 |
| Corning™ Cell Culture Phosphate Buffered Saline (1X) | Thermo Fisher Scientific | Cat#MT21040CV |

| | | |
|---|---|---|
| Bovine Serum Albumin | MilliporeSigma | Cat#A9418 |
| Normal Donkey Serum | Jackson ImmunoResearch | Cat#017-000-121 |
| Normal Goat Serum | Cell Signaling | Cat#5425S |
| Triton-X 100 | MilliporeSigma | Cat#T9284 |
| **Commercial Assays** | | |
| CellTiter-Glo Luminescent Cell Viability Assay | Promega | Cat#G7570 |
| **Experimental Models: Cell Lines** | | |
| VERO C1008 [Vero 76, clone E6, Vero E6] | ATCC | Cat#CRL-158 |
| ARPE-19 | ATCC | Cat#CRL-2302 |
| VK2/E6E7 | ATCC | Cat#CRL-2616 |
| NHEK-Neo – Human Epidermal Keratinocytes, Neonatal, Pooled | Lonza | Cat#00192906 |
| **Software and Algorithms** | | |
| GraphPad Prism 10 | GraphPad | N/A |
| Multi-Point Tool (Cell Counter) | ImageJ | N/A |
| BioRender | BioRender | N/A |

| **COMPOUND DETAILS** | **MOLECULAR FORMULA/SEQUENCE** | **CAS#** | **STRUCTURE** |
|---|---|---|---|
| SM-7368 | $C_{25}H_{22}O_{10}$ | 380623-76-7 | 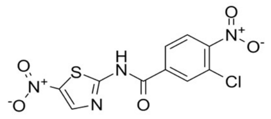 |
| IRAK4-IN-6 | $C_{25}H_{32}N_{10}O_2$ | 2454244-02-9 | 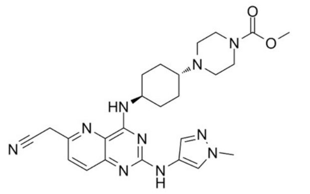 |
| KRAS inhibitor-10 | $C_{30}H_{37}N_3O_5$ | 2578876-75-0 | 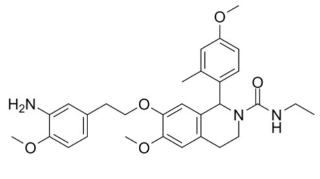 |

| | | | |
|---|---|---|---|
| Niclosamide | $C_{13}H_8Cl_2N_2O_4$ | 50-65-7 | 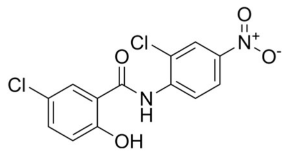 |
| Tecovirimat | $C_{19}H_{15}F_3N_2O_3$ | 869572-92-9 | 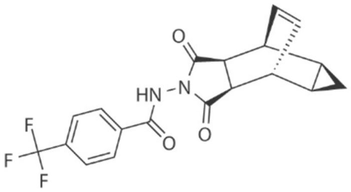 |
